# Quadratic computations maintain neural specificity to natural stimuli across stages of visual processing

**DOI:** 10.1101/2022.11.17.516970

**Authors:** Ryan J. Rowekamp, Tatyana Sharpee

## Abstract

Despite recent successes in machine vision, artificial recognition systems continue to be less robust than biological systems. The brittleness of artificial recognition system has been attributed to the linearity of the core operation that matches inputs to target patterns at each stage of the system. Here we analyze responses of neurons from the visual areas V1, V2, and V4 of the brain using the framework that incorporates quadratic computations into multi-stage models. These quadratic computations make it possible to capture local recurrent computation, and in particular, nonlinear suppressive interactions between visual features. We find that incorporating quadratic computation not only strongly improved predictive power of the resulting model, but also revealed several computation motifs that increased the selectivity of neural responses to natural stimuli. These motifs included the organization of excitatory and suppressive features along mutually exclusive hypotheses about incoming stimuli, such as orthogonal orientations or opposing motion directions. The balance between excitatory and suppressive features was largely maintained across brain regions. These results emphasize the importance and properties of quadratic computations that are necessary for achieving robust object recognition.

## Introduction

Recent advances in computer vision make it possible to achieve human level accuracy in placing visual scenes into categories^1^. Despite these advances, the machine-learning systems exhibit striking vulnerabilities to minor image modifications. For example, image modifications that are imperceptible to human eye can cause the recognition algorithm to make very large errors with high confidence, such as mistaking a school bus for an ostrich^2^. Subsequent machine learning research has been able to trace the origin of such vulnerability to the linearity of the core operation within these networks^3^. Even though these networks apply nonlinearities between stages of processing, at each individual stage inputs are linearly compared to templates. These observations highlight the need to understand nonlinear computational motifs that may be employed by the visual system to achieve fast and accurate visual object recognition.

To address this goal, we have designed a computational model that is scalable and can be used to analyze neural responses across different stages of visual processing. The model extends the standard convolution neural network model to include nonlinearities designed to mimic properties of real neural networks and at the same time remain computationally tractable. Specifically, the model applies quadratic rather than purely linear operation to each patch of the stimulus (or to inputs from the previous stage of processing). A version of this model that combined a quadratic operation at the first stage and a linear convolution at the second stage of the model was previously used to characterize responses of neurons within the visual area V2^4^. Applying this model to responses of neurons from the next stage of processing, the visual area V4, we found that it could not adequately account for these neural responses. Therefore, we developed a model that allows for quadratic matching at each stage of processing, not just the first one (Fig. 1). From a computational perspective, the quadratic operation provides an effective tool for capturing the contributions of local recurrent circuitry at each stage of processing^5^. It can also characterize to multiple overlapping visual features that may affect neural responses locally^6,7^. This includes suppressive features that have been shown to be important for enhancing robustness of neural responses at the primary ^8,9,10,11,12,6^ and the secondary ^13,14,15^ visual areas V1 and V2. Because the quadratic computation is applied at each stage of the model, the framework opens the possibility for discovering the general rules for suppressive interactions.

**Figure 1:**
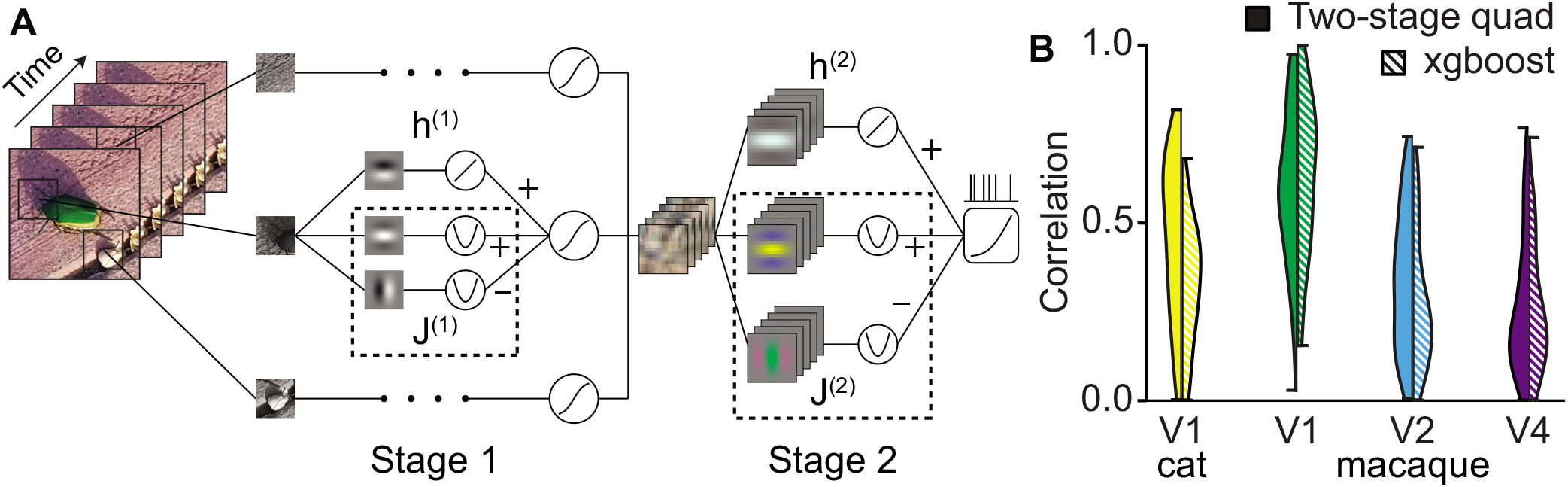
Two-stage quadratic convolutional model accounts for neural responses to natural stimuli across in four brain regions. **A.** Schematic of the model. A colored, spatiotemporal stimulus (left) is broken up into overlapping patches from different times, positions, and colors. These patches are each passed through an identical set of features (middle) to produce a value for each feature at each location. Alternatively, these features are convolved across space, time and color. One of these features is the linear feature while the rest have a quadratic nonlinearity applied to them. The features are combined via a weighted sum and passed through a sigmoid nonlinearity in order to produce a single value at each position. These values are passed through a set of linear and quadratic colored spatiotemporal features (right) before being combined and passed through a softplus rectifier to produce a predicted firing rate. **B.** Comparison with machine learning models. Two-stage quadratic convolutional model performs comparably to non-interpretable models. The quadratic convolutional model has comparable performance with XGBoost, a gradient-boosted decision tree model focused only on predicting responses that is not amenable to interpretation of its parameters (V1: 0.59 ± 0.03 versus 0.63 ± 0.03, V2: 0.34 ± 0.02 versus 0.28 ± 0.02, V4: 0.259 ± 0.015 versus 0.259 ± 0.014).

We use this model to characterize neural responses across different stages of visual processing ^16,17,18^, using data from the primary visual cortex (V1) of the monkey^19^ and the cat^20^, the secondary visual area V2 of the monkey^19^, and finally the area V4 of the monkey^21^. In all cases neural responses were probed with natural stimuli, data from the cat were obtained under anesthesia, while all primate data were obtained from awake animals^19,21^. For each neuron, the fitting procedure adjusted the relative contributions between different stages of processing in the model, the number of features at each stage, and the size patches and pooling masks at the first and second stage. Despite these adjustments, the model remained computationally tractable because it uses at its core the quadratic computational element which by itself allows for convex optimization devoid of local optima^5^.

## Results

### Two-stage quadratic convolutional model matches the performance of machine learning benchmarks

The quadratic convolutional model (Fig. 1) achieved comparable and in some cases substantially better performance than XGBoost (an algorithm that uses random forests to improve performance at the expense of interpretability) ^22^ when accounting for neural responses to novel natural scenes. The model performance was evaluated on a held-out part of the dataset not used for fitting model parameters for a given neuron. In the case of V1 neurons, the quadratic convolutional models actually out-performed the machine learning methods, while reaching > 97% of performance for higher visual processing stages. The high predictive power achieved by the two-stage quadratic convolution model when accounting for neural responses in V1, V2 and V4 validates its use as tool for characterizing the computations performed within these areas.

### Bimodal distribution in the relative dominance of linear and quadratic computations

Classic studies of neurons in the primary visual cortex (V1) divide neurons into two groups, known as the simple and complex cells^23,24,25,26^. These neurons have originally been defined according to the degree to which they summed visual inputs linearly^23,24^. Later studies questioned this classification^27^ arguing that the existing criterion for this classifications^26^ could produce two groups of neurons simply as a result of neuronal thresholds^27^. More generally, one can also view complex and simple cells as a continuum affected by the strength of recurrent contributions ^28^. This raises the question about the coordination between the properties of linear and quadratic features that form subunits of the same neuron. Using the quadratic convolutional framework, we found that neurons had seldomly comparable contributions from linear and quadratic contributions at any stage of their models. Instead, there was a clear dichotomy: at both stages of the model, the variance of the internal state of the subunits was dominated by either the linear or the quadratic contribution but not both at the same time. This reinforces the original definition of simple and complex cells^23,24^, but also generalizes it to apply at the second stage of the model (Fig. 2). Thus, we can define neurons in areas V1, V2, and V4 as belonging at one four classes: linear-linear (LL), linear-quadratic (LQ), quadratic-linear (QL), and quadratic-quadratic (QQ) depending on whether their computations are dominated by linear or quadratic operations at either the first or the second stage of the model.

**Figure 2:**
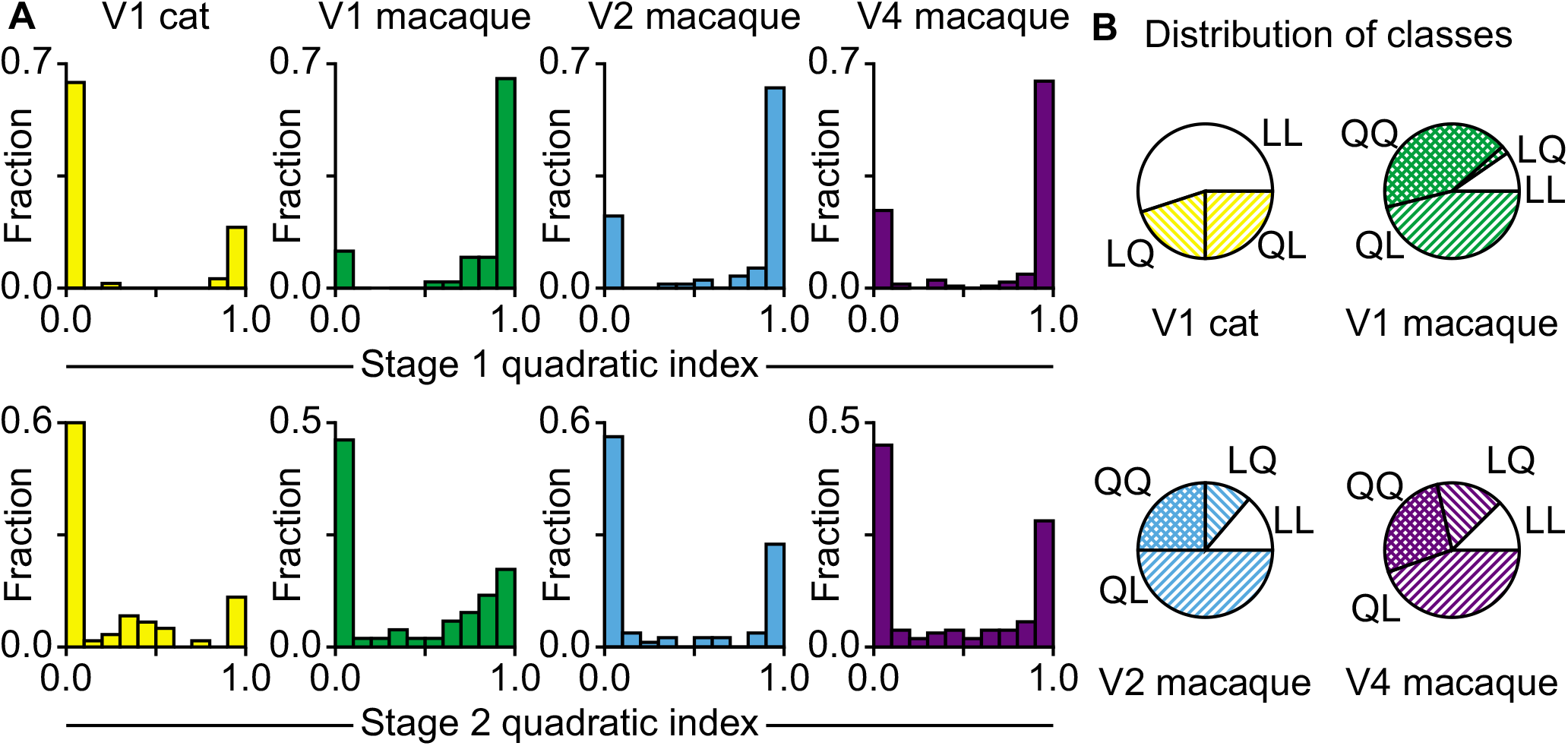
Fraction of variance contributed by quadratic component of first and second stages. **A.** The quadratic index is the variance of the contribution of the quadratic component of a stage divided by the sum of the variances of the linear and quadratic components. Across all areas, most neurons were dominated by either the linear or the quadratic component for both stages, so we divided them into four classes based on which component contributed more to the stages’ outputs. **B.** The distribution of the neurons into four classes: linear in the first layer and linear in the second (LL), linear-quadratic (LQ), quadratic-linear (QL), and quadratic-quadratic (QQ).

The quadratic computations were more common in the primate than in the cat. Neurons with quadratic computations in the first stage of the model comprised 3/4 of the data in area V2 and V4, and a similar but even greater fraction among primate V1 neurons. By comparison, QQ neurons were absent in the cat V1 altogether, with QL neurons comprising 1/4 of the population (Fig. 2). In the primate, the number of quadratic features at the first stage of the model increased for each subsequent brain region along the visual pathway (Fig. S1A, B). Comparing the primate and cat V1, the average number of features at the first stage was not statistically different when counting all cells (both quadratic and linear at the first stage, with linear cells counted as one feature). However, looking at only the quadratic neurons in the first stage, these neurons were less frequent in the cat but had more features than quadratic-at-the-first-stage neurons in the primate (Fig. S1A, B).

Quadratic computations were also important at the second stage, providing a dominant contribution in just under half of neurons in primate V1, V2, V4. In the cat, these neurons represented only 1/4 of the V1 population. Comparing neurons that were quadratic-at-the-second-stage, we found that these neurons had similar number of features across all brain regions studied (no statistically significant differences, (Fig. S1C, ANOVA, *F*[3,129] = 1.69, *p* > 0.05). Because quadratic cells were less common in cat V1, overall across all neurons, the number of features was less in cat V1 compared to V1/V2/V4 of the primate (Fig. S1D, ANOVA, *F*[3, 349] = −4.06, *p* < 0.01; t-test, cat V1 vs macaque V1, *D*(110) = 4.29, *p* < 0.001; vs macaque V2, *D*(138) = −2.80, *p* < 0.01; vs macaque V4, *D*(219) = −3.17, *p* < 0.01). Overall, these results highlight the importance of quadratic computations for multi-stage visual processing. From a computational perspective, one reason for why quadratic computations are important is because they summarize (1) the result of local recurrent computations and (2) make it possible to study interactions between local features, particularly between excitatory and suppressive features. However, before we discuss coordination between excitatory and suppressive features, we describe the main spatial properties that characterize feature selectivity of these visual neurons at the first stage of the model.

### Local selectivity for curved edges

We now describe the main properties of feature selectivity endowed by the first stage of the model, i.e. local feature selectivity. The modeling framework allows the spatial profiles of these features to take arbitrary shapes. Qualitatively, many of the features had edge-like properties, and in some cases curved features, consistent with prior findings from the visual system^29,19,30,31^. To characterize these profiles quantitatively and to connect with prior research, we examined how well these profiles can be explained as combinations of standard feature sets. Four different sets of basis features were used. The first set consisted of standard Gabors^25^. The number of Gabors was equal to the number of local features. The second set consisted from pairs of Gabors, where Gabors in any given pair share all parameters except the spatial phase; the spatial phase differed by 90 degrees between the paired Gabors. The pairs of Gabors represent a standard model for characterizing responses of V1 complex cells^32,25^. The third set of features that we tested where curved Gabors^21^. Our motivation for using the curved feature sets was to describe curvature selectivity that is prominent in area V4^16^ and also observed as early as V1^30,33^. Our final feature set consisted of pairs of curved Gabors, given that such pairing was observed in area V4^21^. We find that sets of independent curved Gabors provided the most accurate fit to the quadratic kernels. This model has the largest number of parameters among the four sets. Nevertheless, the additional parameters could be justified according to the AIC criterion (binomial test, V1 cat: vs individual Gabors *p* < 0.01, vs paired Gabors *p* < 0.01, vs paired curved Gabors *p* < 0.01; V1 macaque: vs individual Gabors *p* < 0.001, vs paired Gabors *p* < 0.001, vs paired curved Gabors *p* < 0.001; V2 macaque: vs individual Gabors *p* < 0.001, vs paired Gabors *p* < 0.001, vs paired curved Gabors *p* < 0.001; V4 macaque: vs individual Gabors *p* > 0.001, vs paired Gabors *p* < 0.001, vs paired curved Gabors *p* < 0.001). Thus, our findings support previous reports of selectivity to curved edges as early as the primary visual cortex ^33,30^.

### Cross-orientation suppression between local features

One of the key features of this modeling framework is that it makes possible to simultaneously estimate excitatory and suppressive features that affect neural responses at each stage of the model. Considering coordination between excitatory and suppressive features we find that in all primate visual areas, from V1 to V4, excitatory and suppressive features had a strong preference for orthogonal orientation at their intersection point (Fig. 4). The bias towards 90° was significant in all primate areas (Kolmogorov-Smirnov test versus uniform distribution; V1 *D*(46) = 0.26, *p* < 0.01; V2 *D*(60) = 0.39, *p* < 0.001; V4 *D*(115) = 0.24, *p* < 0.001). It was not significant in cat V1, but this is most likely due to the small number of neurons with quadratic computation at the local stage (*D*(15) = 0.16, *p* > 0.05).

**Figure 3:**
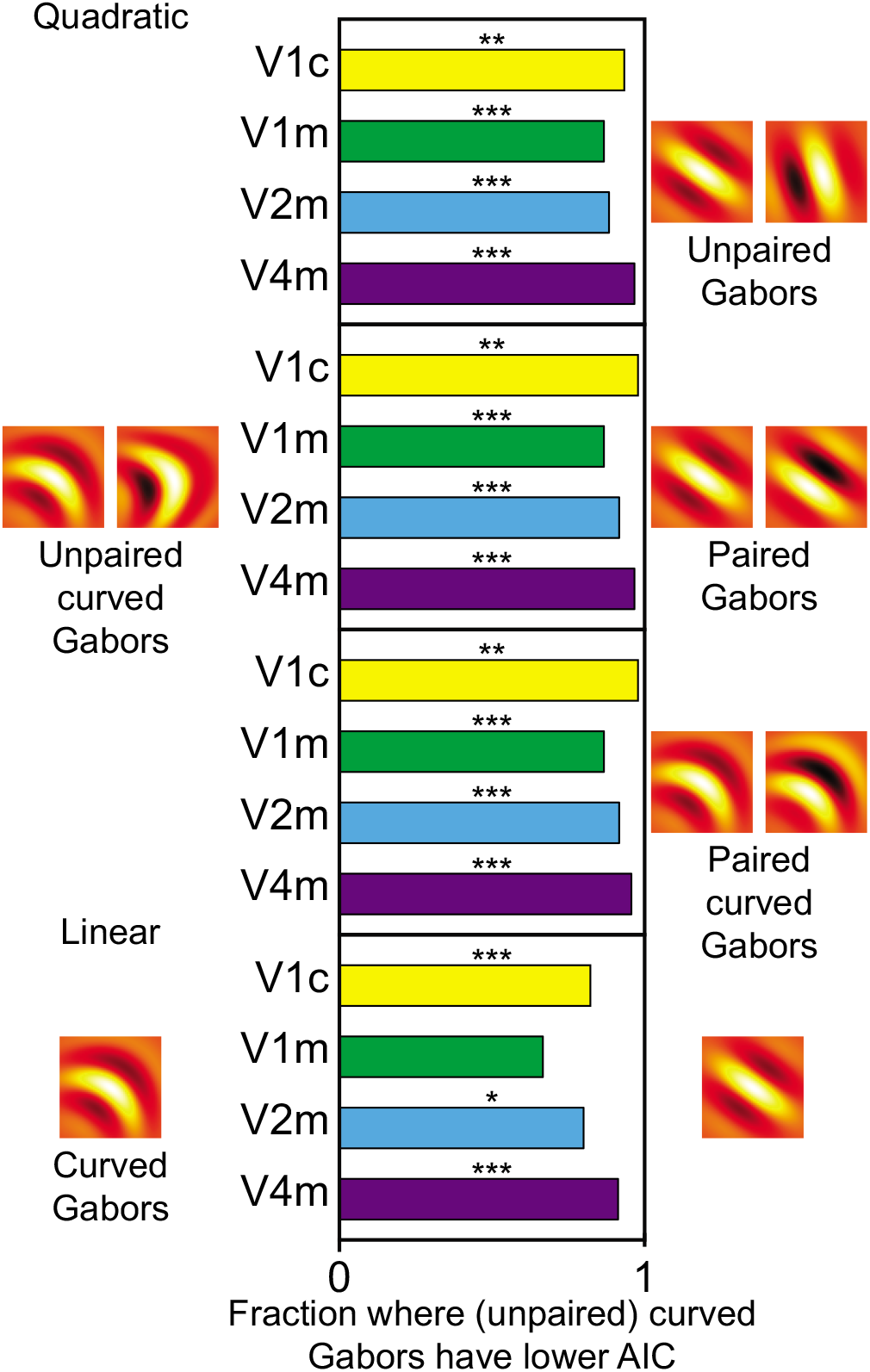
Individual curved Gabor have better AIC scores than alternative models for all areas. Quadratic and linear comparisons only for those cells that were quadratic or linear in the first stage, respectively. Examples of the alternative Gabor models are below their respective bars with the dominant curved Gabor models on the top.

**Figure 4:**
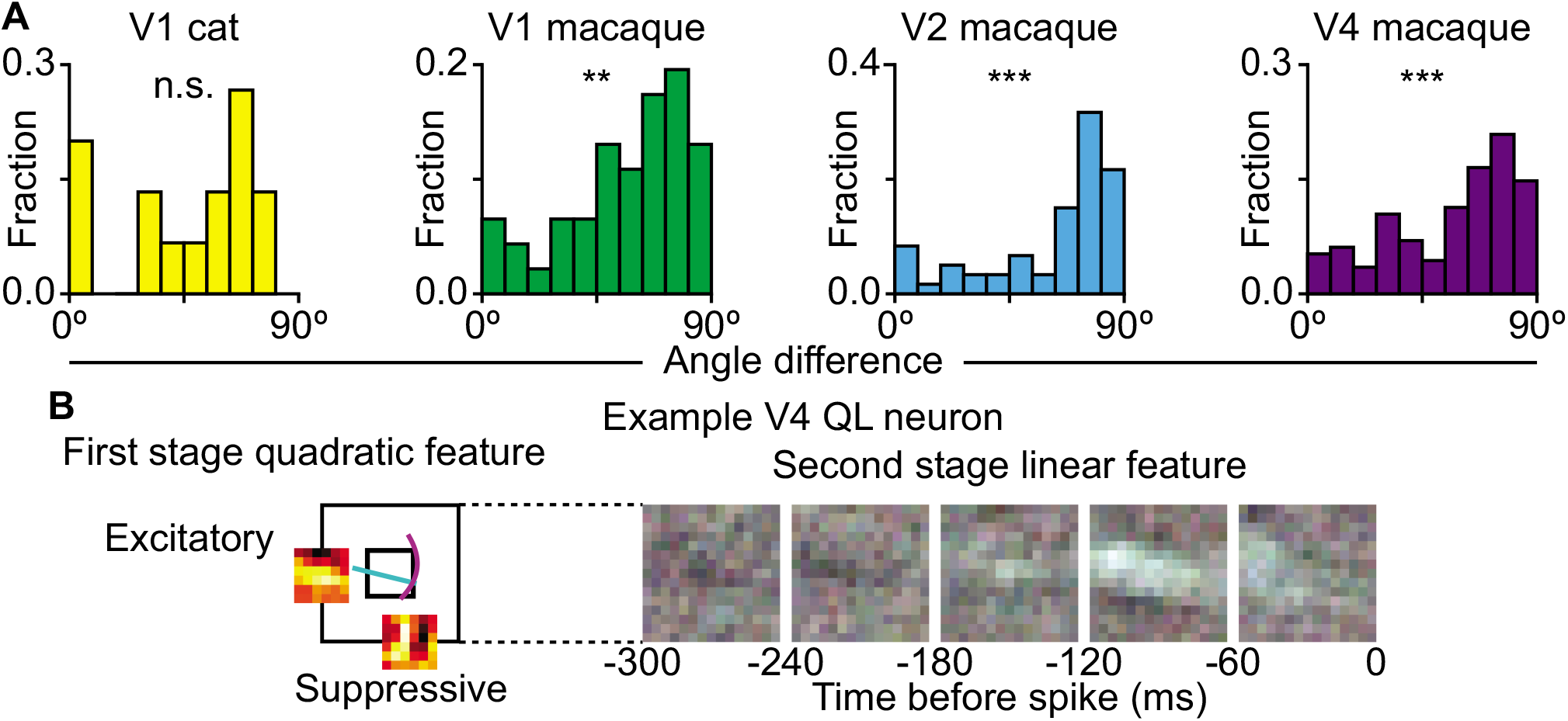
Excitation and suppression are orthogonal to each other in first stage of quadratic cells. **A.** Difference between the mean orientation of the excitatory features and the mean orientation of suppressive features at the position of maximum interaction. The distribution is biased towards orthogonality for macaque V1, V2, and V4 but not cat V1. **B.** An example QL neuron from V4 showing the horizontal preference of the excitatory feature and the vertical preference for the suppressive feature. The inner square is the size of the first stage features and the lines show the peak of the curved Gabors extended out to one *σ* with the horizontal excitatory feature in cyan and the vertical suppressive feature in magenta. Note that they are orthogonal to each other where they intersect.

### Orientation alignment across stages

We now examine the coordination between local and pooling stages of the model. Here we find that the orientations of excitatory features align between stages (Fig. 6A). In other words, the pooling mask has the same primary orientation as the local features it integrates, selecting for continuations of the features preferred by the first stage. This alignment of preferred orientations between local and pooling stages was significant in all areas (Kolmogorov-Smirnov test versus uniform distribution; cat V1 *D*(60) = 0.57, *p* < 0.05; macaque V1 *D*(52) = 0.27, *p* < 0.001; macaque V2 *D*(80) = 0.22, *p* < 0.001; macaque V4 *D*(161) = 0.19, *p* < 0.001). This analysis included both the cases with predominantly linear or quadratic pooling operations at the second stage.

**Figure 5:**
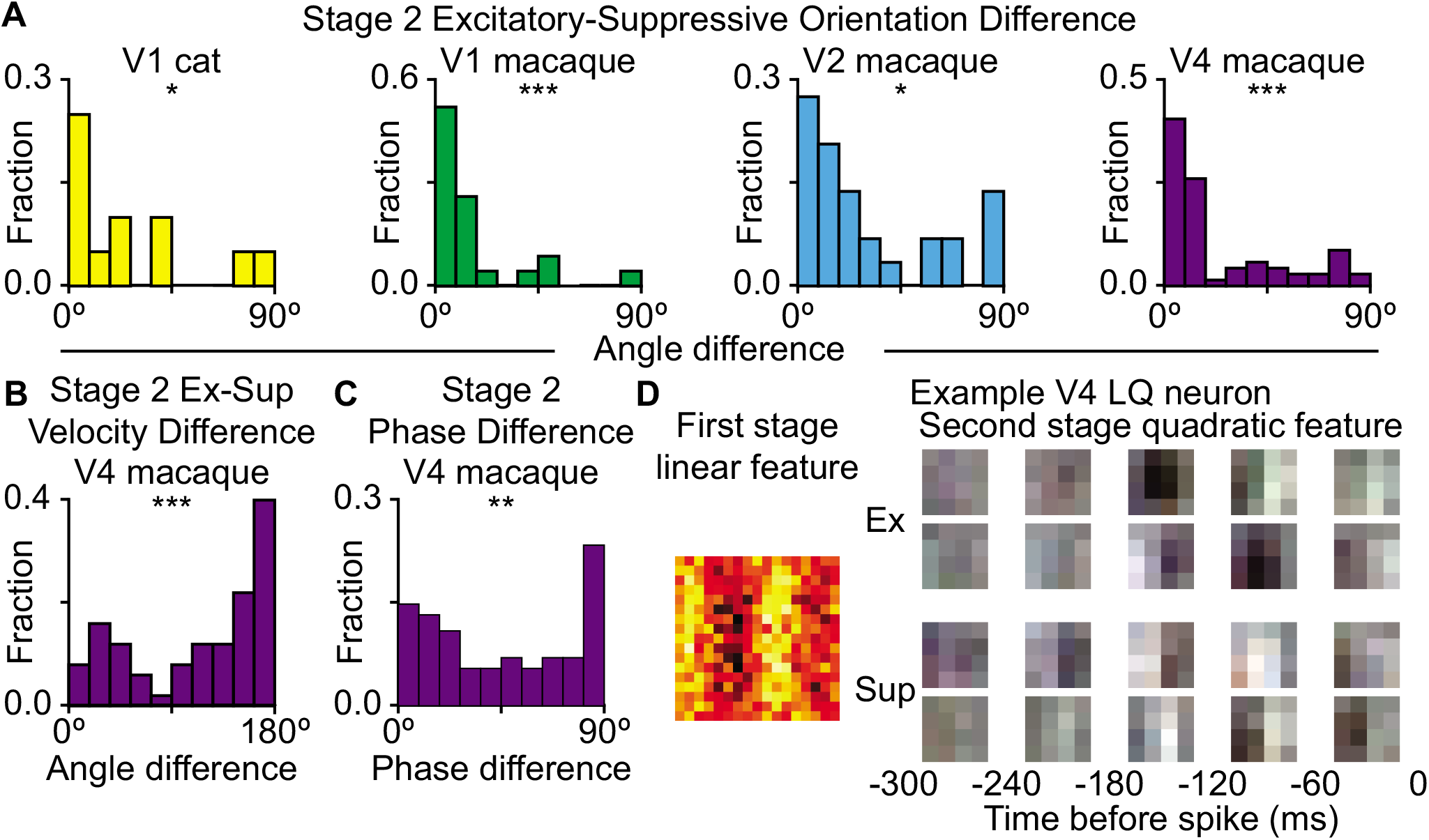
Excitation and suppression are aligned within the second-stage quadratic component. **A.** The dominant orientation of the excitatory and suppressive components of the second stage quadratic cells were aligned for cat V1, macaque V1, V2, and V4. **B.** In macaque V4, second stage quadratic cells excitation and suppression preferred motion of opposite directions. This was not significant for other brain regions (not shown). **C.** For macaque V4 cells quadratic in the second stage, there was a preference for orthogonal phases between the first and second excitatory or suppressive features when there were two or more significant dimensions. This combination of features could provide excitation or suppression to motion that is invariant to the position of the object as it crosses the receptive field. **D.** An example LQ neuron from V4 showing the vertical selectivity in both the excitatory and suppressive feature of the second stage.

**Figure 6:**
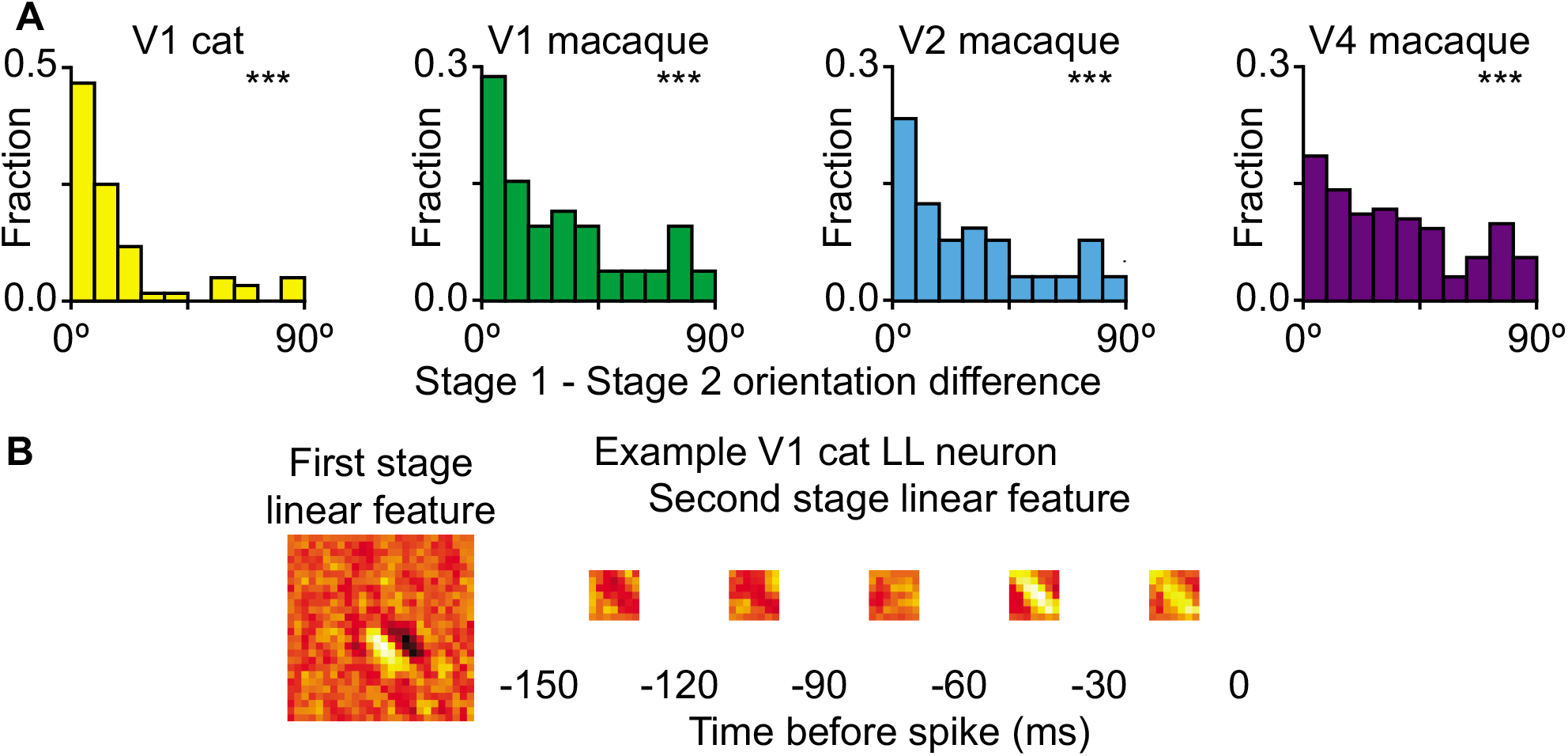
Orientation selectivity aligned between first and second stage. **A.** The difference in the dominant orientation of the first and second stages across V1, V2, and V4. The difference was significantly non-uniform across all three areas (cat V1 *D*(60) = 0.57, *p* < 3 × 10^-19^; macaque V1 *D*(52) = 0.27, *p* < 8 × 10^-4^; macaque V2 *D*(80) = 0.22, *p* < 9 × 10^-4^; macaque V4 *D*(161) = 0.19, *p* < 2 × 10^-5^). **B.** An example cat V1 LL neuron with vertical orientation preference in both its first and second stage features.

### Opponent selectivity to global motion in area V4

Suppressive features were just as prominent at the pooling stage as they were at the first, local stage of the model (Fig. S2). Unlike the cross-orientation suppression that was dominant between local features, here at the pooling stage, for most neurons the excitatory and suppressive features had the same orientation (Fig. 5A; Kolmogorov-Smirnov test versus uniform distribution; cat V1 *D*(12) = 0.39, *p* < 0.05, macaque V1 *D*(23) = 0.64, *p* < 0.001; macaque V2 *D*(29) = 0.38, *p* < 0.001; macaque V4 *D*(69) = 0.47, *p* < 0.001). By construction, the excitatory and suppressive features cannot completely overlap with each other. Therefore, this finding indicates that excitatory and suppressive features differ in other stimulus parameters, such as frequency and/or direction of motion. We did not find systemic differences across the population between the excitatory and suppressive features in earlier brain regions (V1 and V2). However, in area V4 the excitatory and suppressive features were systematically organized along opposing directions of motion (Fig. 5B, Kolmogorov-Smirnov test versus uniform; cat V1 *D*(12) = 0.18, *p* < 0.05; macaque V1 *D*(23) = 0.14, *p* > 0.05; macaque V2 *D*(29) = 0.18, *p* > 0.05; macaque V4 *D*(69) = 0.28, *p* < 0.001). At this point, it is useful to note that selectivity to global motion was stronger in area V4 compared to earlier areas V1 and V2. This was true both for neurons that had predominantly linear or quadratic second stage, (Fig. S3). In area V4, we were also able to detect phase alignment within excitatory and suppressive features in cases where there were at least two significant excitatory or suppressive dimensions (Fig. 5C; Kolmogorov-Smirnov test versus uniform distribution; cat V1 *D*(24) = 0.12, *p* > 0.05; macaque V1 *D*(44) = 0.11, *p* > 0.05; macaque V2 *D*(58) = 0.17, *p* > 0.05; macaque V4 *D*(128) = 0.14, *p* < 0.01). This alignment was not present in earlier stages, possibly because quadratic features were less numerous there (Fig. S1E). In sum, while the balance between excitatory and suppressive features is maintained across brain regions, the quadratic second-stage features were much stronger in area V4 and more clearly organized compared to those in earlier visual areas.

### Coordination between features increases image selectivity

We now examine how the computational motifs described above increase selectivity of neural responses to natural stimuli. First, the overall presence of quadratic computations was crucial to increasing selectivity of neural responses (Fig. 7A-C). For both stages of processing, models that were fit to neural responses without quadratic component produced less sparse responses compared to models where quadratic computations were allowed. This was true in all brain regions, see below for statistical tests, except for cat V1 where quadratics cells constituted a small fraction. The corresponding binomial statistical tests for comparing the full model to the model without quadratic component at either stage were: for cat V1 *p* < 0.001, macaque V1 *p* < 0.001, macaque V2 *p* < 0.001, macaque V4 *p* < 0.001. Comparison of the full model to the model without quadratic component at the first stage yielded: for cat V1 *p* > 0.05, macaque V1 *p* < 0.001, macaque V2 *p* < 0.001, macaque V4 *p* < 0.001. Finally, comparison of the full model to the model without quadratic component at the second stage yielded for cat V1 *p* > 0.05, macaque V1 *p* < 0.001, macaque V2 *p* < 0.001, macaque V4 *p* < 0.001. Therefore, in all primate areas, the quadratic computation at both the first and second stage significantly increased the selectivity of neural responses to natural stimuli.

**Figure 7:**
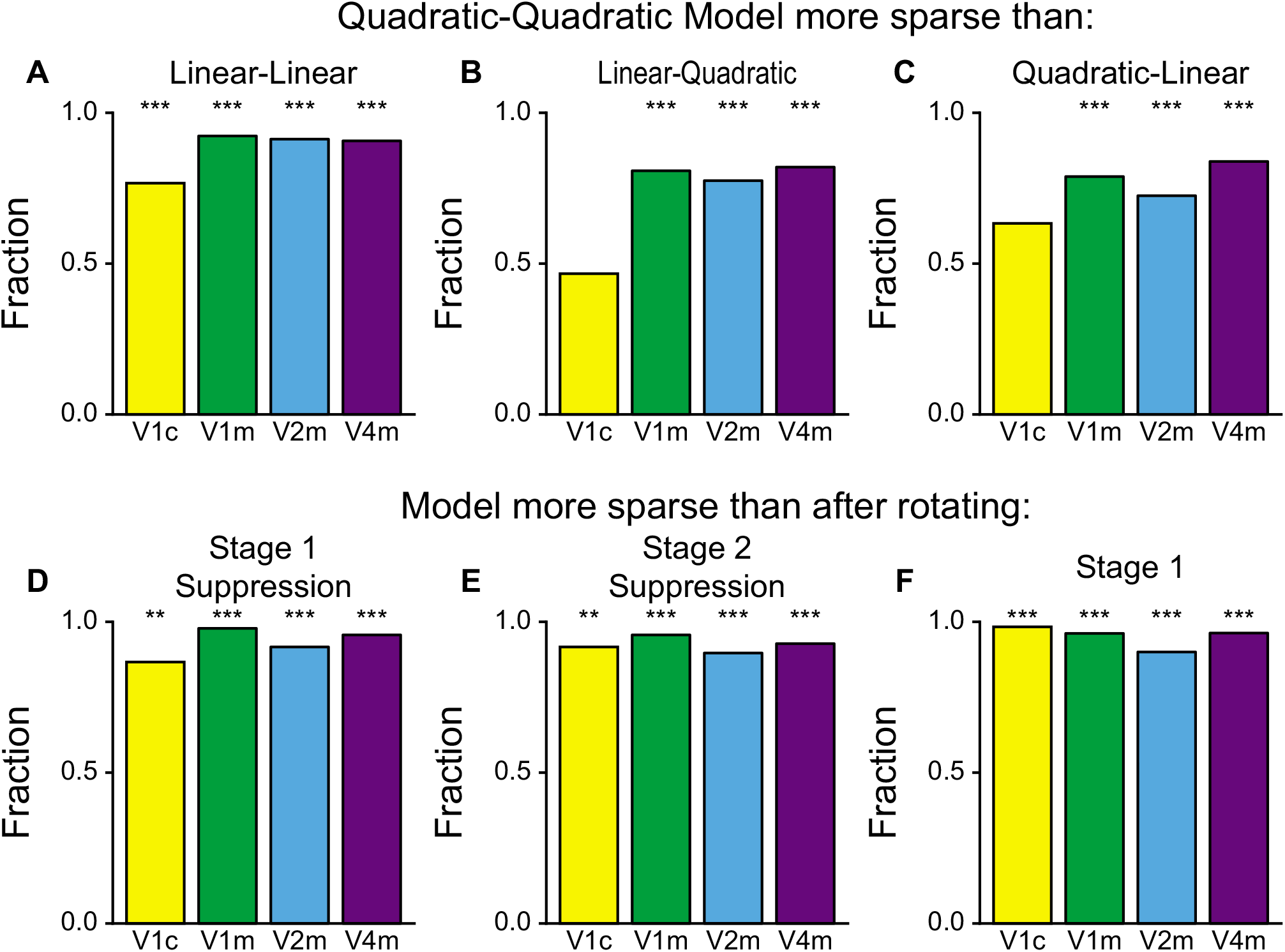
Quadratic computation increase neuronal selectivity to natural stimuli. Alignment between first and second layer, orthogonality of excitation and suppression in the first layer, and alignment between excitation and suppression in the second layer all increase selectivity. For all panels, the bars show the fraction of cells where the responses of the Quadratic-Quadratic model were sparses that the responses of the alterative model. **A-C.** Quadratic-Quadratic model responses were sparser significantly more often than the Linear-Linear, Linear-Quadratic, and Quadratic-Linear alternatives except for cat V1 for the Linear-Quadratic models and Quadratic-Linear models. **D.** For neurons that were quadratic in the first stage, rotating the suppressive features by 90° reduced the sparseness of the modeled responses. **E.** Rotating the suppressive component of the second stage had the same effect on neurons that were quadratic in that stage. **F.** For all neurons, breaking the alignment between the first and second stage by rotating the first stage reduced the sparseness of the responses.

Furthermore, within the quadratic computation, the specific coordination between excitatory and suppressive features was also important for maintaining the selectivity of neural responses. For example, applying a 90° rotation to the suppressive features at either the first or the second stage of the model led to systematic decreases in sparseness across all brain regions studied (Fig. 7D, E). The corresponding statistical tests for rotating the suppression at the first stage decreases the sparseness in cat V1 (binomial test versus 0.5, *p* < 0.01) and in macaque V1 (*p* < 0.001), V2 (*p* < 0.001), and V4 (*p* < 0.001), cf. (Fig. 7D). Similarly, rotating the suppression within quadratic computation at the second stage was significant in all brain regions, with *p* < 0.01 for cat V1, *p* < 0.001 for macaque V1, *p* < 0.001 macaque V2, and *p* < 0.001 for macaque V4, cf. Fig. 7E.

Finally, the alignment of features across stages was also important for maintaining response selectivity (Fig. 7F). Statistically, rotating the first stage relative to the second stage decreased the sparseness in all areas: cat V1 (*p* < 0.001, binomial test), macaque V1 (*p* < 0.001), V2 (*p* < 0.001), and V4 (*p* < 0.001). One way how this alignment can increase selectivity is by selecting for extensions of elongated contours along their preferred direction. It is also worth noting because of correlations present in natural scenes across scales, convolution pre-filtering of visual scenes at the first stage strongly biases the power spectrum of signals received by the second stage (Fig. S4). By aligning with the orientation of the first stage, the second stage is most able to divide the stimuli into different groups and become more selective in its response.

In sum, not only the presence of quadratic kernels was crucial for maintaining the selectivity to natural images, but also the specific coordination between their excitatory and suppressive components was necessary to maintain selectivity. In this regard, it is also important to recall that the balance between excitatory and suppressive quadratic features in stage 1 was maintained at the same level across the four brain regions studied (Fig. S1C, *F*[3, 232] = 2.23, *p* > 0.05).

## Discussion

We have analyzed neural responses across several stages of visual processing in order to determine computational motifs that increase their selectivity to natural inputs. To achieve good predictive power, we found it necessary to extend the standard deep convolutional architecture to allow for quadratic computations at each stage of the models. The quadratic computations not only increased predictive power, but also had specific properties that were essential for maintaining the neural selectivity to natural scenes.

The first unexpected finding was the clear dominance of either the linear or quadratic computation at each stage of the model (Fig. 2). We observed neurons that were linear at the first stage, and quadratic at the second stage, and vice versa, as well as those that were linear/quadratic in both stages. One way to interpret this finding is to note that quadratic computations likely reflect cases with strong local recurrent processing, and in particular in supragranular layers 2/3. Therefore, the observed dichotomy likely reflects the parallel signaling pathways that connect brain areas. One pathway goes from upper cortical layers of the one brain region to the input-recipient layer 4 of the next brain region^34^. The other pathways connect brain regions via thalamus. Taking this anatomy into account, one can interpret a V4 neuron with a strong quadratic computation in both stages of the model as receiving inputs from supragranular layers from two earlier regions. By comparison, a neuron that is predominantly linear at both stages of the model, likely represents pathway that connects input-recipient layers via thalamus. The fact that we also observe cases where quadratic computation at the first stage is followed by a linear computation at the second stage speaks to the crossing between different pathways.

The quadratic features included both excitatory and suppressive features. The suppressive and excitatory features often represented mutually exclusive interpretations of visual inputs. For example, at the first, local stage of the models, excitatory and suppressive features had orthogonal orientation. This finding is consistent with previous analyses of neurons in area V1 and V2 demonstrating that cross-orientation suppression between local features increase the selectivity of these neurons to natural scenes^8,9,10,11,12,4^. Here we extend this finding to other visual areas as well as to other forms of suppressive mechanisms. For example, we find that in area V4, quadratic features at the second stage of the model consists of excitatory and suppressive pairs of features that are selective for opposing directions of global motion. This is interesting because area V4 is not conventionally thought of as motion-selective area^35,36^. Nevertheless, motion signals are known to aid in object segmentation^36^ and that some motion selectivity was detected in V4 neurons by prior studies^37,38^. The quadratic convolutional model shows that selectivity to global motion signals is actually much clearly organized in area V4 compared to earlier areas and is enhanced using similar computational operations as those that increase selectivity to spatial orientation in area V1^8,9,10,11,12^.

For both neural and machine learning systems, the multiple stages of processing are generally believed to yield selectivity to more extended objects^39,30,40^. Despite exhibiting more invariance, real neurons maintain their selectivity across stages of processing^41^. We find that the relative contribution of excitatory and suppressive quadratic features was maintained as at the same level across most the brain regions studied (Fig. S1). The exception was area v1 where suppression was stronger for neurons that had a strong quadratic computation at the second stage of the model. Analysis described above show that quadratic computations contribute to the maintenance of this selectivity. Curiously, in our analysis that increased selectivity came not only from the suppressive computations, but also from the alignment in pooling across stages of processing that also increased response sparseness (Fig. 7F).

Another general phenomenon that is known to take place across the stages of visual processing is the emerging selectivity to more complex, and in particular curved contours ^31,36,16^. At the same time, the selectivity to curved contours can be observed as early as V1 ^30,33^. We did not observe systematic increases in relative curvature of individual features for neurons across stages of visual processing. However, some curvature selectivity could be encoded in the combinations of features, and not in individual features themselves. Supporting this explanation, we find that the number of quadratic features increased from areas V1 to V2 to V4. Our results therefore suggest that the observed selectivity to complex and more curved object contours^31,36^ arises through combinations of quadratic features, with individual features exhibiting some degree of local curvature selectivity. It is also interesting to note that the number of quadratic features was greater in cat V1 compared to primate V1. This finding is consistent with the perspective that more complex computations are carried out at earlier stages of visual processing in smaller brains^42,43^.

Overall, our results point to the importance of quadratic computations for maintaining selectivity and robustness within large hierarchical networks. One of the most important contributions of quadratic computations is that they make it possible to take into account excitatory and suppressive features. The magnitude of excitatory and suppressive contributions to the quadratic was balanced across the brain regions. Across subsequent brain areas, the excitatory and suppressive features are aligned to represent mutually exclusive hypotheses about the natural world. This specific alignment increased selectivity. Overall, these results emphasize the importance and properties of quadratic computations that are necessary for achieving robust object recognition.

## Methods

### Electrophysiological recordings

We fit our model to neuronal recordings from four different datasets. The first dataset was from the primary visual cortex of cats^20^. Tetrode electrodes recorded spike trains as the anesthetized animals were exposed to natural movie stimuli. The movies were 128 × 128 pixels with a resolution of 0.12° per pixel. We downsampled the movies to 32 × 32 pixels before fitting our model.

The second dataset was recorded from the primary visual cortex of macaque monkeys^44^. Tungsten electrodes recorded the spike trains while the animals fixated on the screen for juice rewards. The movies were natural image sequences or natural images with simulated saccades. The size of the movies was 50 × 50 pixels with a resolution of 0.1-0.5° per pixel. We took the central 40 pixels and downsampled them to 20 × 20 pixels before fitting.

The third dataset was recorded from visual area V2 of macaque monkeys^19^. The spike trains were recorded while the animals performed a fixation task for liquid reward. The stimuli were random patches of images chosen to favor high contrast and were scaled to match the size of the measured classical receptive field. We downsampled the stimuli to 20 pixels.

The final dataset was recorded from visual area V4 in awake macaque monkeys^21^. The spikes were also recorded during a fixation task. during a fixation task. The stimuli were natural movies displayed at a 120 × 120 pixe resoluation with a fixed size of 14 × 14°. The movies were downsampled to 20 × 20 pixels and were analyzed in color.

### Low-Rank Quadratic Convolutional Model

The model seeks to make a prediction (*Ŷ_t_*) of a neuron’s response *Y_t_* (measured in action potentials/spikes per time bin) given the stimulus **X**_*t*_ presented.

The stimulus begins with a size of 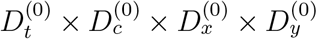, where *D* is the size along the time, color, horizontal, and vertical directions. In the first layer, the stimulus is convolved with *H*^(1)^, *U*^(1)^, and *V*^(1)^. *H*^(1)^ is the linear term with a single component. *U*^(1)^ and *V*^(1)^ combine to form the quadratic term and have *m*^(1)^ components. Each component has size 1 × 1 × *d*^(1)^ × *d*^(1)^. The outputs of these convolutions are

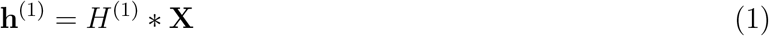

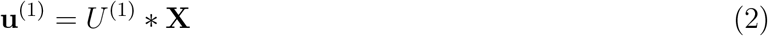

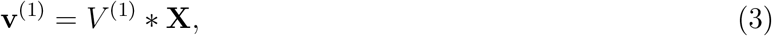

and they are combined like

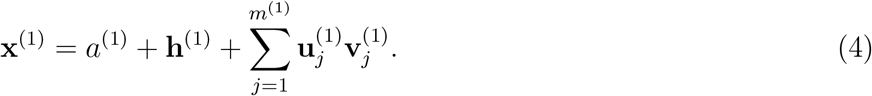

*a*^(1)^ is a scalar bias. This is passed through a logistic function to give the output of the first layer:

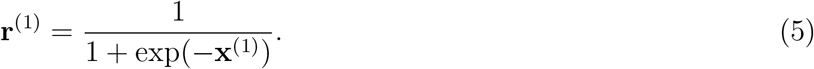

The second layer is similar to the first with the following exceptions: the kernels are dotted with the input rather than convolved, the components have size 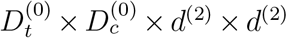 where 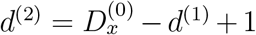, and the activation function is

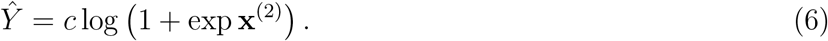

*c* is a positive scalar that adjusts the mean response. In total, the model has nine sets of parameters: *a*^(1)^,

### Fitting the model

The model was implemented using the Keras python package using Theano as a backend. We used the adadelta optimizer and terminated optimization when the Poisson log-likelihood on the validation set failed to improve for more than two passes.

The model has nine hyper parameters: the spatial size of the first layer kernels (*d*^(1)^), the ranks of the quadratic filters for each layer(*m*^(1)^, *m*^(2)^), individual regularization parameters for each layer’s linear and quadratic filters (*λ*_*H*^(1)^_, *λ*_*H*^(2)^_, *λ*_*UV*^(1)^_, *λ*_*UV*^(2)^_), and regularization parameters for the difference *UU^T^* – *VV^T^* for each layer (*ϵ*_*UV*^(1)^_, *ϵ*_*UV*^(2)^_). In order to choose the values of the hyperparameters, we used the Spearmint package^45^, which models the performance in the space of hyperparameters as a Gaussian process and suggests new values that have the highest expected improvement. For each set of hyperparameters, we fit the model four times using different random divisions of the data. The seeds for these random divisions were the same for each set of hyperparameters. The performance of hyperparameter sets was evaluated using the mean correlation between the predictions of the models and the observed responses on their respective validation sets. We ran 100 sets of hyperparameters and selected the set with the best correlation for further analysis. Analysis of the parameters averaged them across the four models while analysis of the performance averaged their predicted firing rates.

### Benchmark model

In order to determine how well our model was at predicting neural responses, we also fit XGBoost models ^22^.

### Evaluating model performance

To calculate the performance of the models, we evaluated

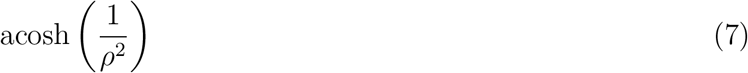

for subsets of the test dataset of varying lengths (where *ρ* was the correlation) and calculated a linear regression versus the inverse of the length. The bias term of this regression is the estimated value for infinite data, and inverting Eq. 7 provides our estimate of the correlation between our model predictions and the observed responses.

### Dividing cells into linear and quadratic classes

We calculated the standard deviations of the linear (**x_h_**) and quadratic 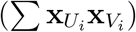 contributions to each later to get *σ*_**h**^(1)^_, *σ*_*UV*^(1)^_, *σ*_**h**^(2)^_, and *σ*_*UV*^(2)^_. With these we calculated the fraction of the layer’s variance contributed by the quadratic component

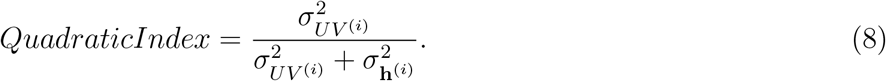

If the quadratic index was greater than 0.5, we classified the layer as quadratic (Q1 for the first layer and Q2 for the second). Otherwise, we classified it as linear (L1 or L2). Each layer was classified independently, creating four classes: Linear-Linear (LL), Linear-Quadratic (LQ), Quadratic-Linear (QL), and Quadratic-Quadratic (QQ).

### Feature fitting using differential evolution

In order to extract perceptually relevant properties from the parameter weights of the model, we fit the kernels as various types of features using differential evolution^46^, which randomly initializes sets of parameters and generates new sets by taking a set and modifying random values using the values of the other sets. If the child set has a lower error than its parent, it takes its parent’s place. The specific variation of the algorithm is termed “rand/2/bin” ^4^.

The types of features that we used to analyze each neuron depended on what class (LL, QL, LQ, QQ) the neuron belonged to. For neurons dominated by the quadratic term on the first stage, we fit the quadratic kernel *J*^(1)^ using curved Gabor wavelets:

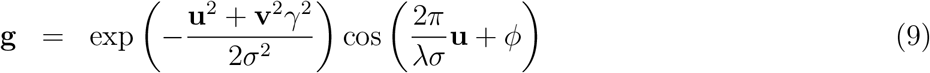

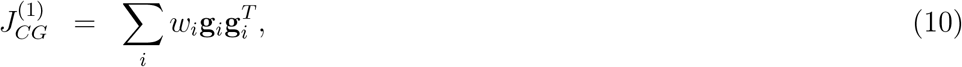

where

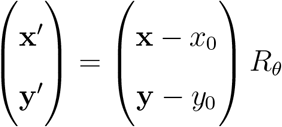

and

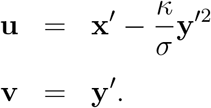

The parameters for each curved Gabor wavelet are the weight (*w*), the position (*x*_0_, *y*_0_), the orientation (*θ*), the curvature (*κ*), the aspect ratio (*γ*), the size (*σ*), the spatial wavelength (*λ*), and the phase (*ϕ*).

Variations were also tested. Uncurved models fixed *κ* to be zero while paired models had pairs of Gabor wavelets with identical parameters except that *ϕ* was set to be 0 and *π*/2. The combination of paired and unpaired, curved and uncurved created four alternative models.

For the second stage quadratic cells, we fit two moving sign functions:

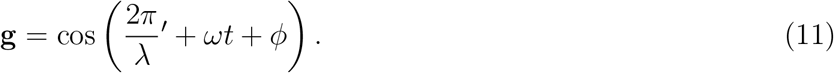

Figure 1A shows the structure of the model. The stimulus has two-spatial dimensions, a color dimension (in the case of V4 data), and a temporal dimension. A series of linear spatial kernels are convolved across each position, time, and color (if applicable). One of these kernels is treated linearly while the rest are passed through a quadratic function before being combined into a single value at each part of the stimulus. These values are passed through a sigmoid function to create the input for the second layer. Another set of linear and quadratic kernels are applied to this input to create a single value that is passed through a rectifying nonlinearity to produce a predicted firing rate.

### Local orientation difference

Starting with the parameters of a curved Gabor, one can calculate the local orientation using

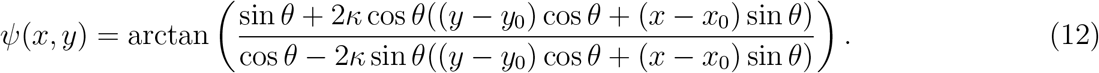

Each local orientation was given a weight with three factors: |*w*| (the Gabor’s weighting in *J*), the weight of the curved Gaussian envelope *a*(*x,y*), and the norm of the projection of the Gabor into the significant eigenvectors

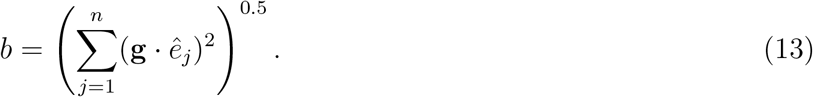

Using the weighted angular mean

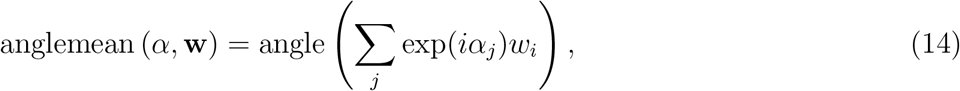

we calculated the mean excitatory or suppressive orientation at a location as

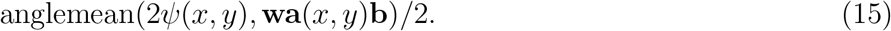

The factors of two causes *ψ* and *ψ*+*π* to add together rather than cancel each other out since the orientation is not directional.

We selected the point of maximum interaction between excitation and suppression by summing the weights at each location for the excitatory and suppressive Gabors respectively and then multiplied them together. Using the absolute angular difference

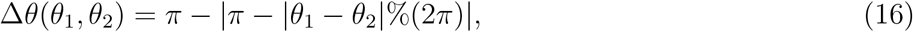

the difference between the excitatory and suppressive orientation preference is

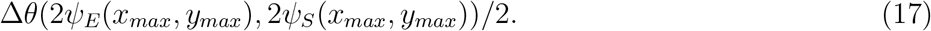

Once again, the factor of two makes the calculation direction independent.

### Measuring orientation and direction selectivity in second-layer features

Given a spatiotemporal feature *a_x,y,t_*, we calculated the dominant orientation by first calculating the Fourier transform with respect to the two spatial dimensions: *â_l,m,t_*. We also calculated the orientation of each Fourier component

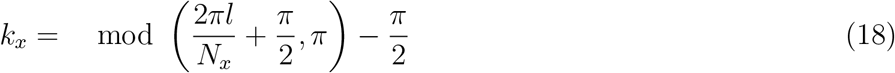

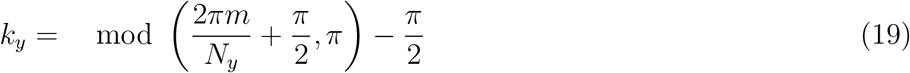

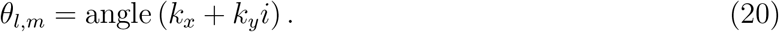

These angles are averaged together weighted by the Fourier transform

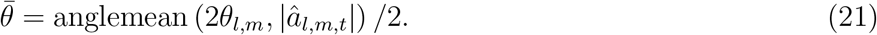

The weights for the zero spatial frequency components and the Nyquist frequency components are set to zero since the orientation is undefined. The factors of two ensures that *θ* and *θ* + *π* are treated as the same orientation.

Measuring the direction is done similarly with a few modifications. The Fourier transform is taken with respect to the temporal dimension in addition to the spatial dimensions to produce *â_l,m,n_*. The angle is also calculated slightly differently:

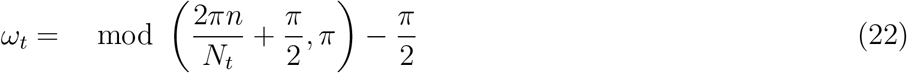

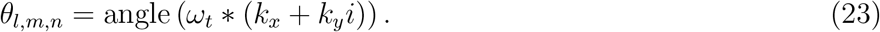

The angles are averaged using

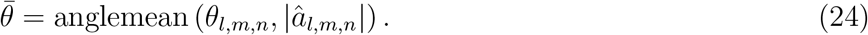

In addition to the non-oriented components zeroed above, the components for *ω_t_* = 0 were also zeroed as they had no directed motion. The factors of two are removed from this variation because *θ* and *θ* + *π* are no longer equivalent.

---

**Figure S1:**
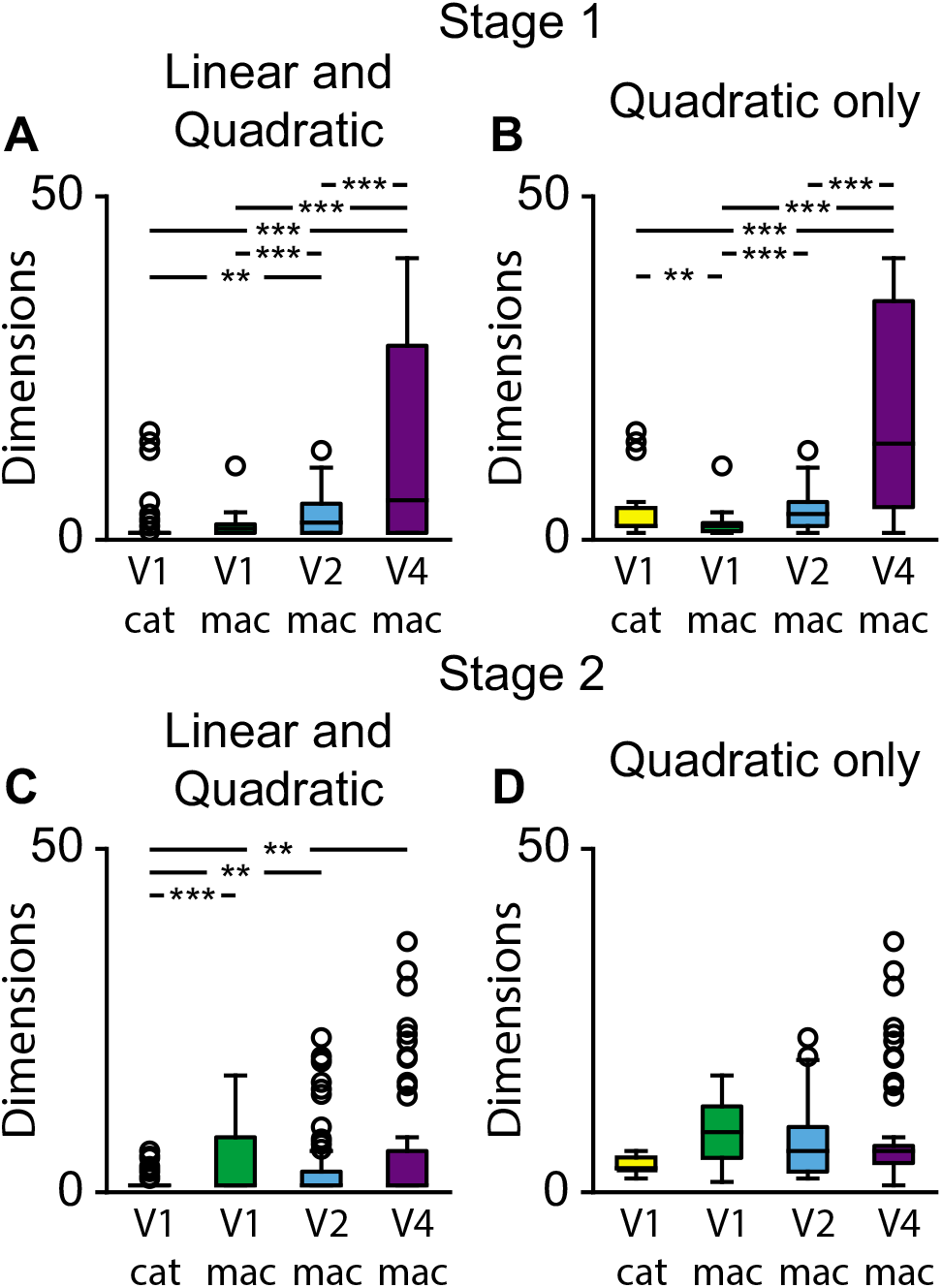
V4 neurons have more dimensions in the first stage. **A, B.** The number of dimensions in the first stage for the quadratic cells only or for all cells counting the linear cells as having one dimension. For both measures, V4 has more features than earlier visual areas. **C, D.** The number of dimensions for the second stage. The number of dimensions was not significantly different across areas except that cat V1 had significantly fewer when linear cells were included.

**Figure S2:**
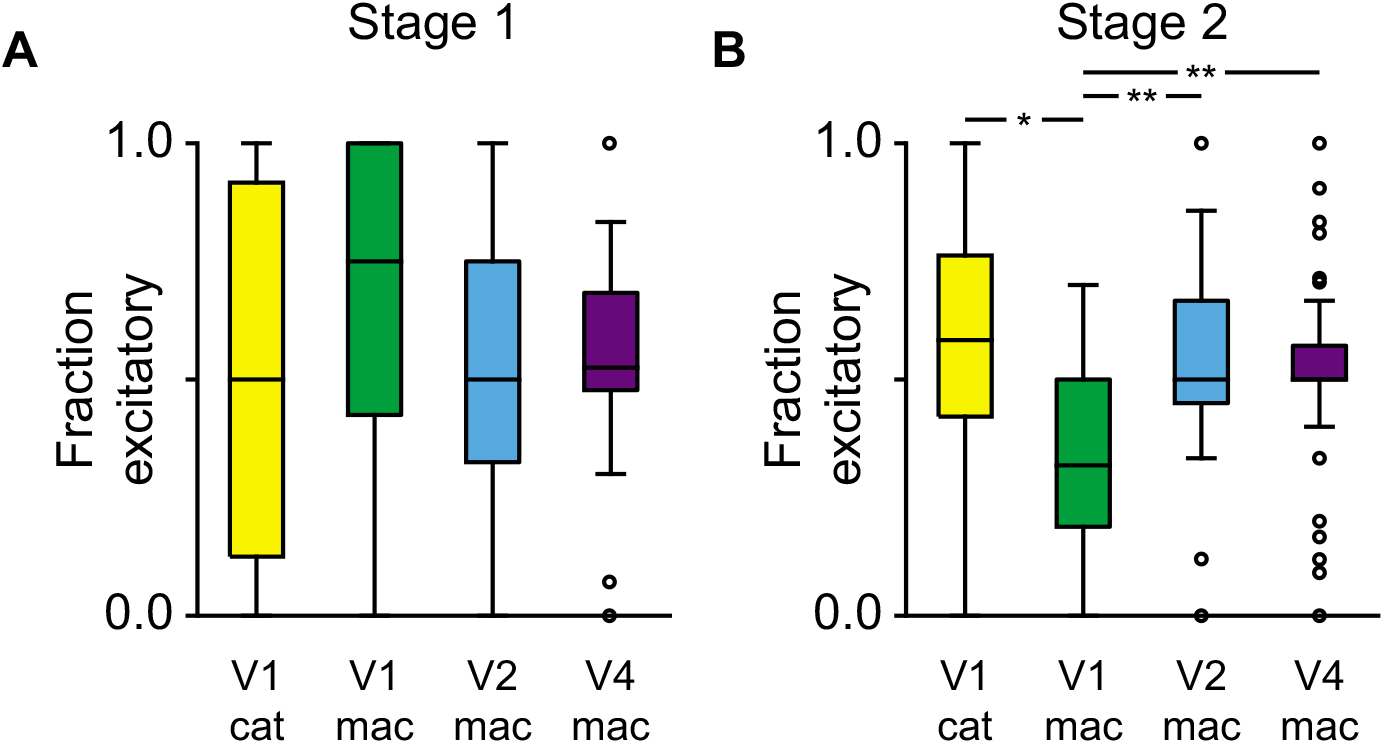
Balance of excitation and suppression across brain areas. **A.** The distribution of the relative strength of excitation in the first stage as a fraction of the total eigenvalue weights. There were no significant differences between the areas. **B.** The excitatory fraction for the second stage. Macaque V1 had more suppression than the other areas.

**Figure S3:**
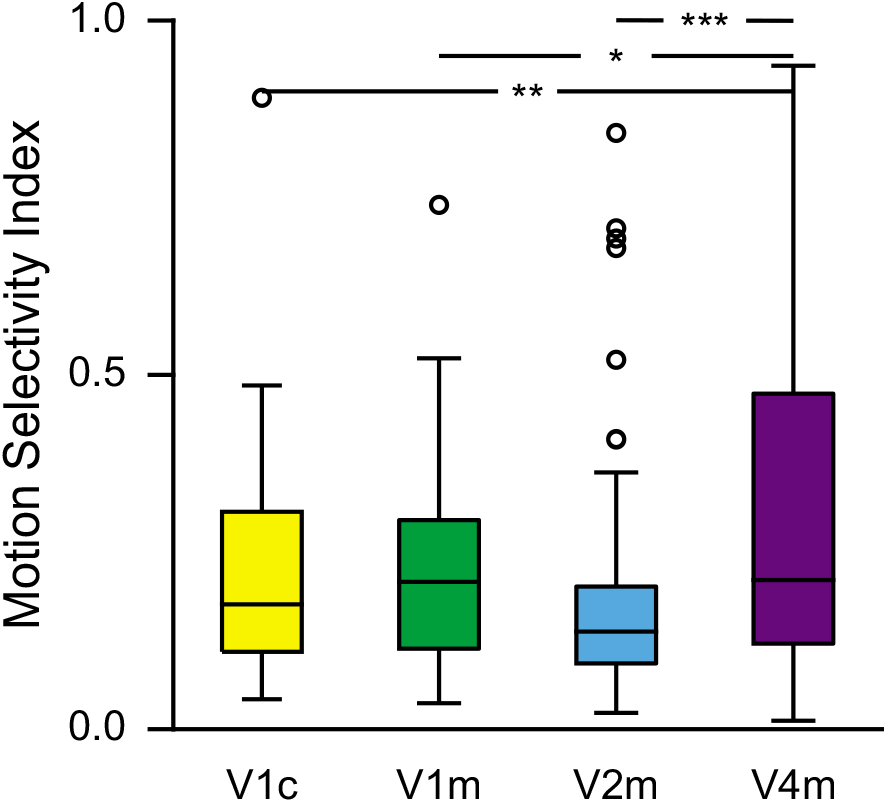
V4 cells are more selective for motion. The dominant feature of the second stage of V4 neurons had a higher motion selectivity index than the other areas we studied, i.e. the features were less separable into spatial and temporal components. This indicates that V4 has a higher capacity to select for motion.

**Figure S4:**
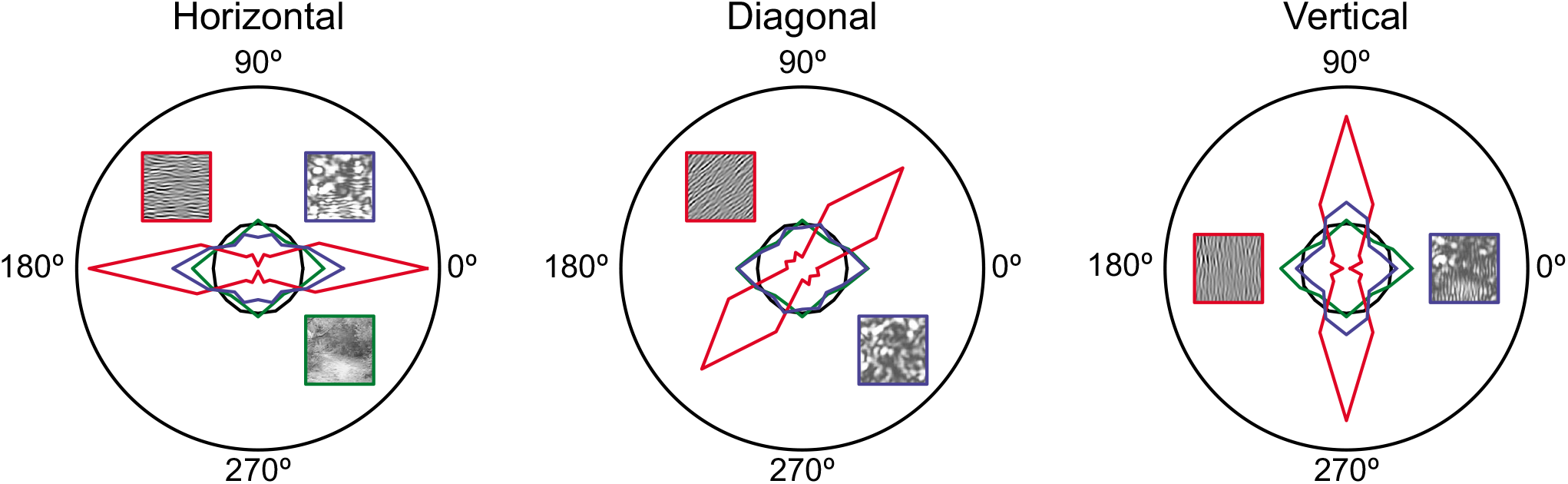
Orientation distribution at the first stage biases inputs to the second stage towards its orientation. Distribution of oriented energy in gaussian noise (black), natural movies (green), the first stage with a linear feature (red), and the first stage with a quadrature pair of excitatory features. The insets show examples of the associated images. The distributions for horizontal, diagonal (45°), and vertical features are shown. The linear example created the largest shift in the distribution relative to the natural image inputs, but the quadratic example also shifted the distribution towards the preferred direction of the first stage’s features.

